# Establishment of a Replicon System for Bourbon Virus and Identification of Small Molecules that Efficiently Inhibit Virus Replication

**DOI:** 10.1101/2020.04.24.058693

**Authors:** Siyuan Hao, Kang Ning, Xiaomei Wang, Jianke Wang, Fang Cheng, Safder S. Ganaie, John E. Tavis, Jianming Qiu

## Abstract

Bourbon virus (BRBV) was first isolated from a patient hospitalized at the University of Kansas Hospital in 2014. Since then, several deaths have been reported to be caused by BRBV infection in the Midwest and Southern United States. BRBV is a tick-borne virus that is widely carried by lone star ticks. It belongs to genus *Thogotovirus* of the *Orthomyxoviridae* family. Currently, there are no treatments or vaccines available for BRBV or thogotovirus infection caused diseases. In this study, we reconstituted a replicon reporter system, composed of plasmids expressing the RNA-dependent RNA polymerase (RdRP) complex (PA, PB1 and PB2), nucleocapsid (NP) protein, and a reporter gene flanked by the 3’ and 5’ UTR of the envelope glycoprotein (GP) genome segment. By using the luciferase reporter, we screened a few small molecule compounds of anti-endonuclease that inhibited the nicking activity by parvovirus B19 (B19V) NS1, as well as FDA-approved drugs targeting the RdRP of influenza virus. Our results demonstrated that myricetin, and an anti-B19V NS1 nicking inhibitor, efficiently inhibited the RdRP activity of BRBV and virus replication. The IC_50_ and EC_50_ of myricetin are 2.22 μM and 4.6 μM, respectively, in cells. Myricetin had minimal cytotoxicity in cells, and therefore the therapeutic index of the compound is high. In conclusion, the BRBV replicon system is a useful tool to study viral RNA replication and to develop antivirals, and myricetin may hold promise in treatment of BRBV infected patients.

## 1. Introduction

Bourbon virus (BRBV) is a member in the genus *Thogotovirus* of the *Orthomyxoviridae* family. There are seven genera in the family of *Orthomyxoviridae*, which are segmented negative-strand RNA viruses (Kawaoka & Palese, 2006), including four types of influenza viruses (influenza virus A, B, C, and D), thogotovirus, quaranjavirus, and isavirus. Many viruses in the *Orthomyxoviridae* family are important pathogens to humans or animals. The epidemic/pandemic influenza A viruses have caused millions of human deaths in the past century and are still circulating, posing a huge threat to human health and the economy. Thogotoviruses mainly circulate in domestic animals, such as sheep, cattle, and camels, causing neural diseases and abortion. Two thogotoviruses, thogoto virus and dhori virus, have been reported to infect humans and cause deaths (Butenko, et al., 1987; Kosoy, et al., 2015). Human antibodies against thogoto virus and dhori virus have been detected in Europe, Asia, and Africa (Hubalek & Rudolf, 2012; Filipe, et al., 1985). It has been reported that thogoto virus can cause human infections in a vector-free manner, possibly by an aerosol route (Butenko, et al., 1987), highlighting the potential to infect humans in a large population.

BRBV was first isolated from a blood sample of a hospitalized male patient from Bourbon County, Kansas, USA, in the spring of 2014 at the University of Kansas Hospital (Kosoy, et al., 2015). The patient died due to a complex syndrome of leukopenia, lymphopenia, thrombocytopenia, hyponatremia, and increased levels of aspartate aminotransferase and alanine aminotransferase. Due to the high level of viremia and unique identification of the virus in the serum taken from the patient, BRBV was believed to be the cause of the illness and the death of the patient. Later, two additional cases of BRBV infection have been reported. In 2015, a patient from Payne County, Oklahoma, USA tested positive for neutralization antibodies to BRBV before fully recovering (Savage, et al., 2017). In June 2017, a 58-year-old woman from Missouri died from an infection of BRBV after she had been misdiagnosed for a significant period of time (Bricker, et al., 2019). Not surprisingly, a high seroprevalence of BRBV–neutralizing antibodies in raccoons (50%) and white-tailed deer (86%) was detected in Missouri, USA (Jackson, et al., 2019). However, the Center for Disease Control and Prevention (CDC), USA, currently does not know if the virus may be found in other areas of the United States, since a seroprevalence of BRBV infection has not been evaluated outside of these epidemic regions. Nevertheless, BRBV is the first species of the genus *Thogotovirus* of the *Orthomyxoviridae* family to be identified as a human pathogen in the New World (Lambert, et al., 2015).

BRBV is widely carried by the lone star tick (*Amblyomma americanum*), a species that is aggressive, feeds on humans, and is widely distributed across the East, Southeast, and Midwest States. BRBV was found in 3 pools of lone star ticks from retrospective tests in 39,096 ticks from northwestern Missouri, 240 km from Bourbon County, Kansas, USA (Savage, et al., 2017). The human BRBV and the strain isolated from tick pools share >99.0% sequence at the amino acid level and 95.0% identity at the RNA sequence level (Savage, et al., 2017). BRBV replicates in cell lines derived from the hard ticks *Amblyomma*, *Hyalomma* and *Rhipicephalus* (Lambert, et al., 2015). Together with the geographic location of the BRBV infection and the geographic distribution of a number of *Amblyomma americanum* ticks (Savage, et al., 2017; Savage, et al., 2018), these studies strongly suggest that the lone star tick is a vector of BRBV transmission to humans.

Little is known about the biology of BRBV. Due to the high mortality of BRBV infection, specific treatments with antiviral drugs are in demand to save lives. In this study, we established a replicon reporter of BRBV. We then used the replicon reporter to examine anti-influenza drugs that target viral RNA-dependent RNA polymerase (RdRP) and endonuclease inhibitors of human parvovirus B19 (B19V) for inhibition of BRBV RdRP activity and BRBV replication in HEK293T and Vero cells.

## 2. Materials and Methods

### 2.1. Cells and Virus

#### Cell cultures

HEK293T cells (ATCC CRL-11268) and Vero cells (gifted from Dr. Maria Kalamvoki) were cultured in Dulbecco’s modified Eagle’s medium (DMEM) (HyClone, catalog no. SH30022.01; GE Healthcare Life Sciences, Logan, UT) supplemented with 10% fetal bovine serum (FBS; catalog no. F0926, Sigma, St. Louis, MO) at 37°C under a 5% CO_2_ atmosphere.

#### Virus

BRBV-KS (#NR-50132) was obtained through BEI Resources, NIAID, NIH. The virus was amplified on Vero cells once, aliquoted, and stored at −80°C in the Hemenway 4037 BSL3 Lab of the University of Kansas Medical Center and used for all subsequent studies. A biosafety protocol to work on the virus in the BSL3 Lab was approved by the Institutional Biosafety Committee (IBC) of the University of Kansas Medical Center.

### 2.2. Compounds

T705 (Favipiravir) and VX-787 were purchased from AdooQ Bioscience (Irvine, CA). Baloxavir (S-033447; Baloxavir acid) was purchased from MedChemExpress (Monmouth Junction, NJ). Flavonoid compounds used in this study were commercially acquired as follows: #7 (Idofine #2030), #9 (Sigma, #70050), #135 (AldrichSelect #361173301), #201 (Sigma, # S0327), #860 (Enamine, #Z1918018629). They all had a purity of ≥95%. All compounds were dissolved in DMSO (Sigma, St. Louis, MO) as stock solutions at 10 mM, and stored at −80°C.

### 2.3. Virus titration assays

#### Plaque assays

Cells were seeded in six-well plates at a density of 2 ×10^6^ cells and were confluent the following day. We used the cell growth media to serially dilute the virus stock at 10-fold. 200 μl of the diluent were added to each well and incubated for 1 h with gently rocking the plate every 20 min. After removing the virus diluent, 2 ml of overlay media, 1% methylcellulose (Sigma, M0387) in DMEM with 5% FBS, were added to each well. The plates were incubated at 37°C under 5% CO_2_ for 5 days. After removing the methylcellulose overlays, cells were fixed using the 10% formaldehyde solution for at least 30 min and stained with 1% crystal violet solution.

#### Reverse transcription and quantitative PCR (RT-qPCR)

Viral RNA was isolated from virus preparation or supernatants of virus-infected cells using a viral RNA extraction kit (Quick-RNA Viral Kit, R1035, Zymo Research) by following the manufacturer’s instructions. M-MLV reverse transcriptase (#M368A, Promega) was used to reversely transcribe viral RNA with the reverse PCR primer according to the manufacturer’s instructions. Then, the transcribed cDNA was subjected to qPCR using primers and a TaqMan probe targeting to the NP gene of BRBV (**Table 1**) with a 7500 Fast Real-Time PCR System (Applied Biosystems). In order to eliminate the contamination of free viral RNA, we quantified the nuclease digestion-resistant viral genome copies (vgc) numbers. To this end, 100 μl of virus preparations were treated with 25 units of Benzonase (Sigma) for 30 min before RNA isolation.

**Table 1.**
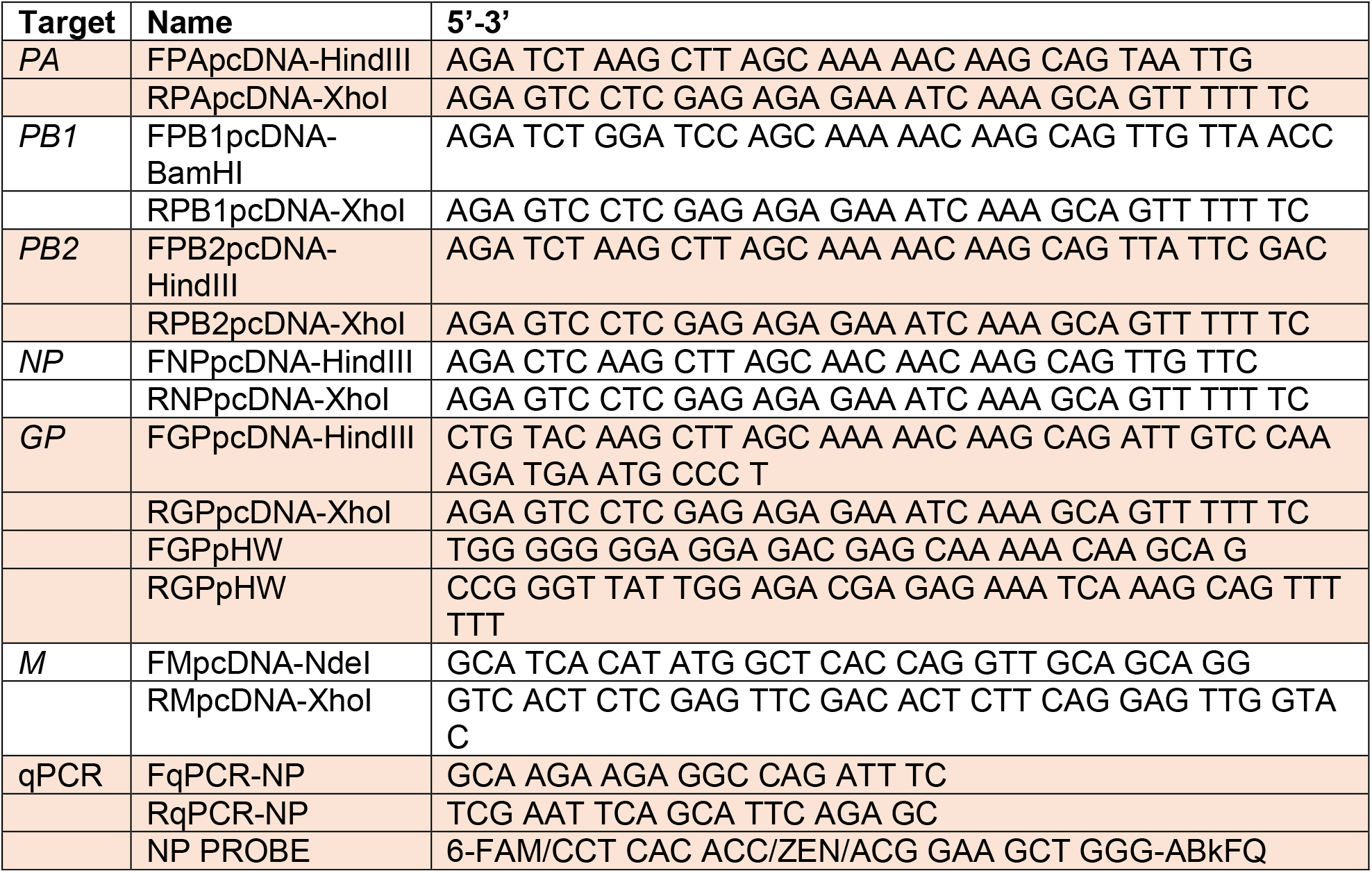
Primers and probe used in the study.

### 2.4. Viral cDNA cloning and plasmid constructions

#### pcDNA3-based viral cDNA plasmids

BRBV RNA was isolated from a stock of BRBV-KS passaged on Vero cells. cDNA of each viral RNA genome fragment was synthesized using the reverse primer and amplified by PCR. The amplification program started with one cycle at 48°C for 45 min and one cycle at 94°C for 2 min. These cycles were followed by 40 cycles at 94°C for 20 sec, 52°C for 30 sec, and 72°C for 40 sec; the program ended with one cycle at 72°C for 5 min. The PCR products were visualized by agarose gel electrophoresis, extracted, and cloned into pcDNA3 vector (Invitrogen), resulting in pcDNA-(BRBV) PA, PB1, PB2, M, NP, and GP plasmids.

#### pHW2000-GP plasmid

The plasmid pHW2000 (Hoffmann, et al., 2002) contains the human ribosome RNA polymerase I (pol I) promoter and the murine terminator sequence separated by two BsmBI sites. The pol I promoter and terminator elements are flanked by a truncated immediate–early promoter of cytomegalovirus (CMV) and by the bovine growth hormone polyadenylation signal (bGHpA). The pcDNA3-GP was used as a template to amplify the GP cDNA with a primer set that has the BsmBI sites (**Table 1**). After digestion of the PCR products with BsmBI, the fragments were cloned into the pHW2000 vector, which resulted in the pHW2000-(BRBV)GP plasmid.

#### Reporter plasmids

pHW2000-ΔCMV-GP was constructed by deletion of the CMV promoter in pHW2000-GP. Then, we constructed pHW-ΔCMV-GP-GFP and pHW-ΔCMV-GP-gLuc plasmids inserting the GFP ORF through the NdeI and Eag I sites of the GP ORF and replacing the GP ORF with the *Gaussia luciferase* (*gLuc*) ORF, respectively, into the pHW2000-ΔCMV-GP plasmid (**Fig. 4A**).

The sequences of all the primers are listed in **Table 1**. All the constructed plasmids were sequenced to confirm the cloned cDNA sequences at MCLAB (South San Francisco, CA). DNA sequence analyses were performed using SnapGene 4.0 (SnapGene, Chicago, IL).

### 2.5. Plasmid DNA transfection

HEK293T cells were transfected using the PEImax (Cat# 24765-2, MW 40,000, Polyscience, Inc.) at a ratio of 1:3 of DNA: PEI in 200 μl of opti-MEM (Invitrogen) (Wang, et al., 2018). The total amounts of plasmid DNA were kept constant (2-3 μg per well of 6-well plate) in each group by supplementation with an empty vector.

### 2.6. BRBV reporter assays

#### BRBV GFP reporter assay

pHW-ΔCMV-GP-GFP was co-transfected with the four pcDNA-PA, PB1, PB2, and NP plasmids into HEK293T cells confluent in six-well plates. After 7 days post-transfection, the GFP signal was observed under a fluorescent microscope (Nikon Ti-S). The transfection of pHW-ΔCMV-GP-GFP with three pcDNA plasmids (PB1, PB2, NP) was set as a negative control.

#### BRBV luciferase reporter assay

pHW-ΔCMV-GP-gLuc and the four pcDNA-based plasmids were co-transfected into HEK293T cells in 96-well plate. At 3 days post-transfection, 10 μl of the cell culture supernatants were taken and mixed with 50 μl of Working solution or luciferase activity, using the *Gaussia* Luciferase Flash Assay kit (Pierce™, #16159). Luminescent signals were detected on a Synergy H1 microplate reader (BioTek, Winooski, VT). Three pcDNA plasmids (PB1, PB2, and NP) and pHW-ΔCMV-GP-gLuc co-transfected cells were set up as a negative control.

### 2.7. Half cytotoxic concentration (CC_50_) assay

The cytotoxicity of compounds on HEK293T and Vero cells was evaluated using the CytoTox-Glo™ Cytotoxicity Assay kit (Promega, #G9290), according to the manufacturer’s instructions. Cells were seeded at 4 × 10^4^ cells per well in 96-well plates. After overnight incubation, different concentrations of the compounds diluted in culture media were added to each well. Cells treated with DMSO were used for the negative control. Cell viability was measured after three days of incubation.

### 2.8. Half maximal effective concentration (EC_50_) assay

Confluent monolayers of Vero and HEK293T cells in 96-well plates were inoculated with 10 pfu of BRBV. Different concentrations of the compounds, diluted in medium, were added to the wells. Each concentration of the compounds was tested in duplicate per experiment, and the experiment was repeated three times. At 3 days post-infection, culture supernatants were collected and used to quantify the nuclease digestion-resistant vgc numbers using RT-qPCR.

### 2.9. Western blotting

Cells were collected by centrifugation and lysed using a 1 ml tuberculin syringes. The cell lysates were loaded, along with 2.5 μl of a pre-stained protein ladder (#P008; GoldBio, St. Louis, MO), and separated on an SDS-polyacrylamide gel. Proteins were transferred onto a polyvinylidene difluoride membrane (#IPVH00010; Millipore, Bedford, MA). The membrane was blocked and probed with primary and secondary antibodies sequentially. Signals were visualized by enhanced chemiluminescence, and the pre-stained protein ladder was imaged under bright light simultaneously under a FujiLAS4000 imaging system.

An anti-BRBV M protein polyclonal antibody was produced in rats by immunizing them with the BRBV M protein, which was expressed and purified in bacteria, following a protocol previously described (Sun, et al., 2009). Anti-β-actin (#A5441) antibody was purchased from Sigma (St. Louis, MO).

### 3.0. Statistical analysis

Calculations of IC_50_, CC_50_ and EC_50_ and statistical analysis were done by using GraphPad Prism version 8.0. Error bars represent means and standard deviations (SD), and statistical significance (P value) was determined by using the Student t test.

## 3. Results

### 3.1. BRBV-KS infection of HEK293T cells

We propagated BRBV-KS, obtained from BEI Resources (www.beiresources.org/), in Vero cells. At 7 days post-infection, when most of the cells appeared cytopathic, the media were collected and centrifugated. The supernatant was collected, aliquoted, and stored at −80°C. The virus was titrated on Vero cells with a titer of 1.5 × 10^8^ plaque forming units (pfu)/ml (**Fig. 1A**). We next developed a RT-qPCR to quantify the viral genome copy (vgc) numbers in the BRBV stock. We found a high correlation of the vgc numbers with the pfu titrated (γ=0.9875) (**Fig. 1B**). In the following experiments, we used the RT-qPCR to quantify the virus.

**Fig 1.**
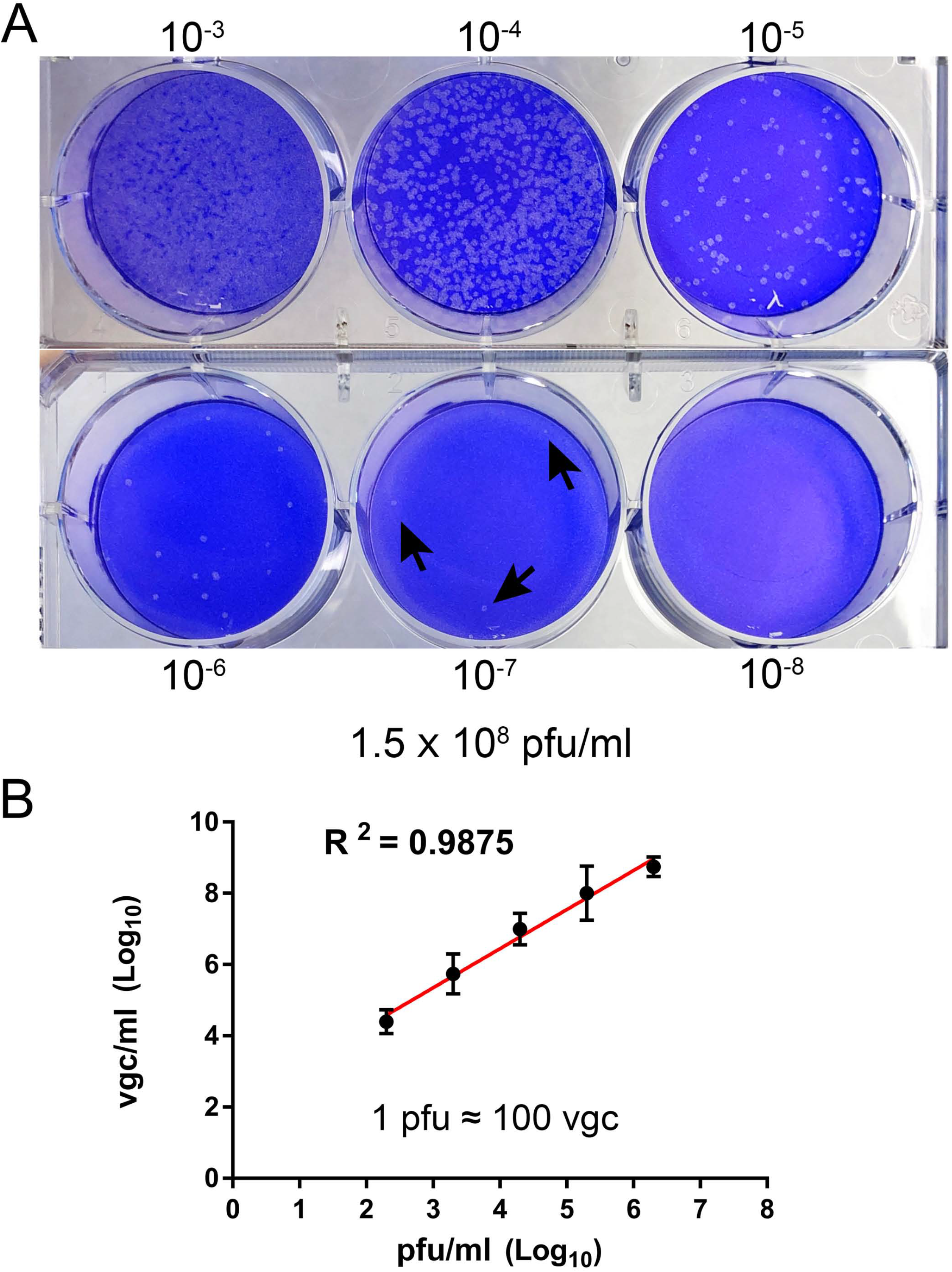
Titration of BRBV stock. **(A) Plaque assay.** Vero cell monolayers on a 6-well plate were infected with 200 μl of serial 10-fold diluted BRBV-KS as indicated. Plaques were visualized by staining with crystal violet after 5 days of infection on Vero cells. Arrows indicate plaques. **(B) Comparison between plaque assay and RT-qPCR.** The same virus stock was diluted by serial 10-fold dilution. RT-qPCR was used to quantify the viral genome copy (vgc) number. The linear regression has a R^2^ of 0.9875, indicating 1 plaque forming unit (pfu) equals ~100 vgc. The data presented are averages and standard deviations from three independent experiments, with each experiment analyzing samples in duplicate.

We tested human HEK293T cells for infection of BRBV for comparison with infection in Vero cells. Virus growth was determined over the course of infection, and infected cells were collected at 6 days post-infection for Western blotting of matrix (M) protein expression. The results showed that BRBV infected HEK293T cells displayed similar growth kinetics as those in Vero cells (**Fig. 2A**), which had a peak of virus production of approximately (~) 10^9^ vgc/ml in the media, as well as detection of the M protein at a size of ~30 kDa (**Fig. 2B**).

**Fig 2.**
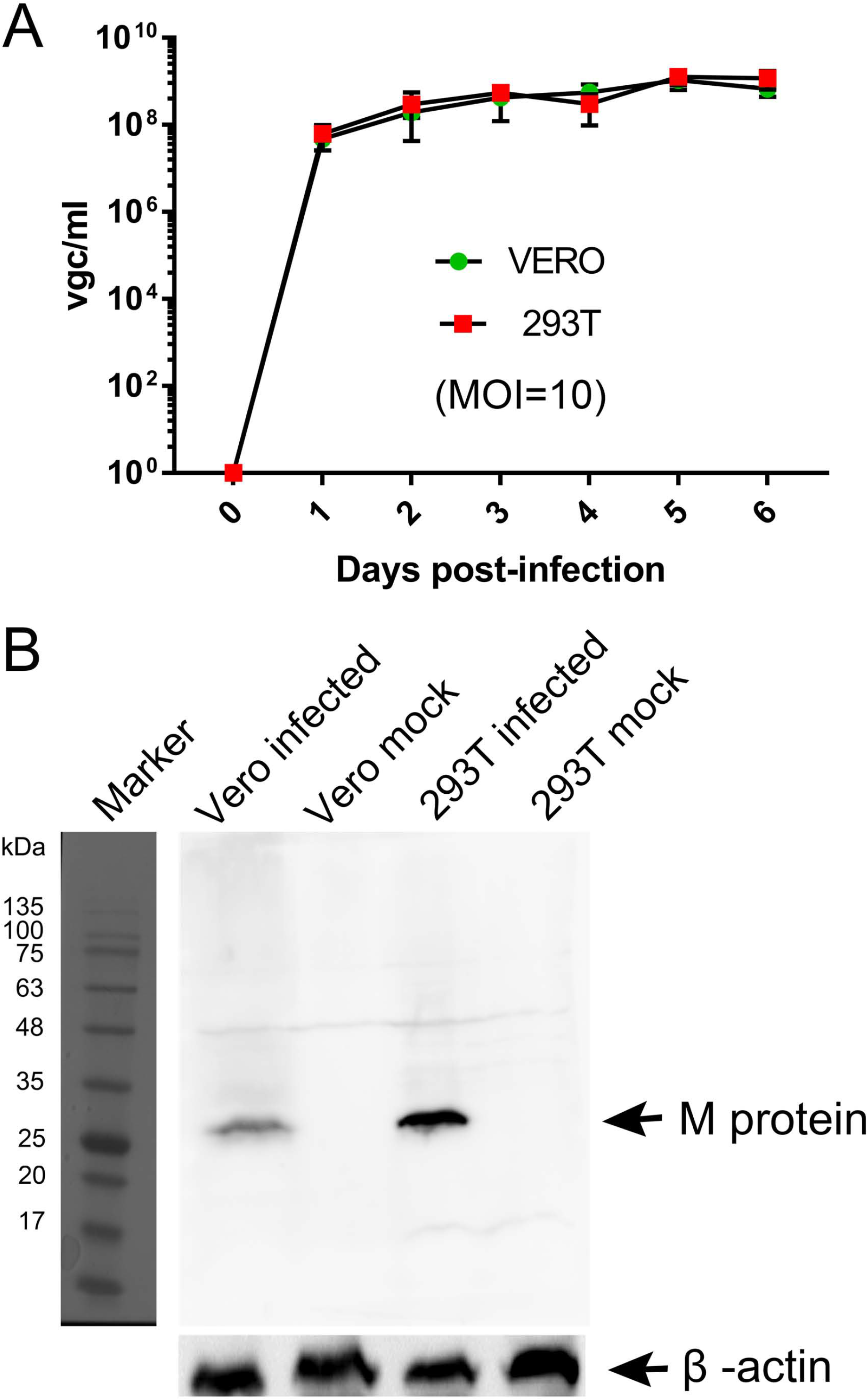
Comparisons of BRBV-KS infection between Vero and HEK293T cells. **(A) BRBV growth kinetics.** Cells were infected with BRBV-KS at an MOI=10 pfu/cell. Cell culture media (supernatants) were collected daily for 6 days. Viral RNA was extracted from the samples of supernatants and then titrated by a RT-qPCR assay. The data presented the means with standard deviations, which are obtained from three independent experiments, with each experiment analyzing samples in duplicate. **(B) Western blotting.** The M protein in BRBV-infected Vero and HEK293T cells was detected at ~30 kDa as indicated, but not in mock-infected cells. β-action was probed as a loading control.

### 3.2. Cloning of the BRBV cDNA fragments

To clone the 6 cDNAs of the 6 genome segments of BRBV-KS, we designed primers complementary to the 3’ UTR and 5’ UTR of the originally published genome sequences of BRBV-KS, KU708255 (*GP*), KU708254 (*PB1*), KU708253 (*PB2*), KP657749 (*NP*), KP657750 (*M*), and KP657748 (PA) (**Table 1**). Each viral gene was reversely transcribed using each forward primer and PCR amplified using respective paired primers (**Table 1**). The PCR cDNA fragments were first cloned into pcDNA3 and sequenced. Sequences of *GP*, *PA*, and *PB2* genes were confirmed as identical to those originally deposited in GenBank; whereas *NP*, *PB1* and *M* genes have 10, 7 and 1 nucleotide variations, respectively (**Fig. 3A**), which result in amino acid variations in NP and PB1 (**Fig. 3B**).

**Fig 3.**
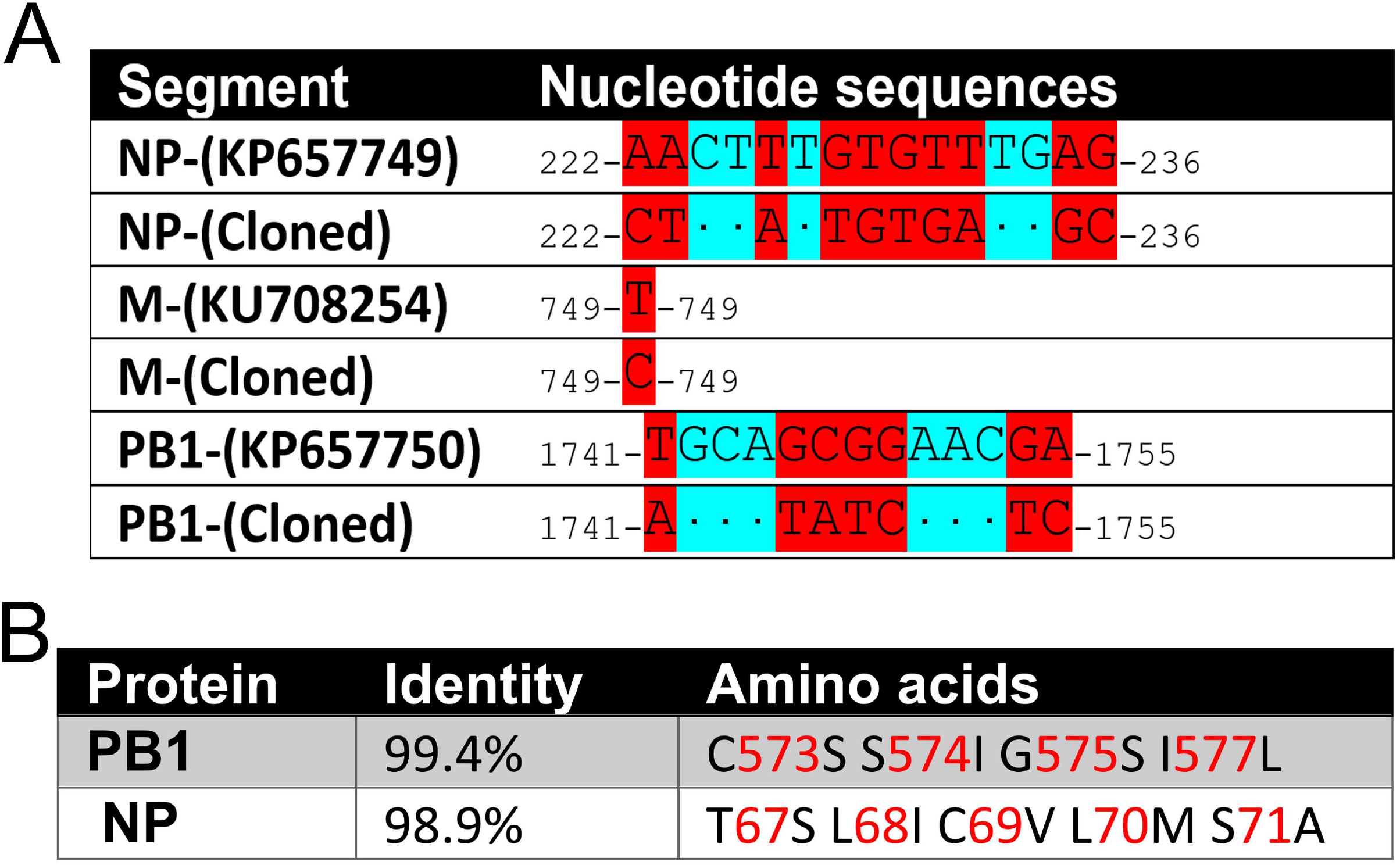
Sequence variations of the cloned BRBV genome segments. **(A) Nucleotide sequence variations.** Nucleotide changes in BRBV-KS in comparison to the sequences of BRBV genome segments deposited in Genbank. **(B) Amino acid sequence variations.** Percent identity at the amino acid level and amino acid alternations of the PB1 and NP proteins between sequences of BRBV-KS and the ones deposited in GenBank. No variations were found in PA, PB2, and GP proteins.

We next used the set of pHW primers (**Table 1**) that had the BsmBI sites to amplify the GP cDNA and cloned it in a BsmBI-digested pHW2000 bidirectional cloning vector (Hoffmann, et al., 2002). The pHW2000-GP clone that had the right direction of BRBV cDNAs was confirmed by sequencing, in which there is the immediate early promoter of CMV, RNA polymerase II (pol II) promoter, a pol I terminator in the 3’ untranslated region (UTR), a ribosome RNA pol I promoter and a bGHpA in the 5’ UTR.

### 3.3 Establishment of a viral RNA dependent RNA polymerase (RdRP) activity assay, a replicon reporter system of BRBV

Working on BRBV requires a biology safety level 3 (BSL3) facility, thus, a replicon system of BRBV is important for assessing antivirals in a BSL2 setting and for high throughput screening of antivirals. Based on the pHW2000-GP plasmid, we inserted a GFP open reading frame (ORF) and a secreted *Gaussia luciferase* (*gLuc*) ORF between the 3’ and 5’ UTR of the GP segment (**Fig. 4A**). When pHW2000-GFP was co-transfected with pcDNA-PA, PB1, PB2, and NP plasmids in HEK293T cells, at 7 days post-transfection, green fluorescence was clearly observed in cells transfected with pcDNA-PA, PB1, PB2, and NP plasmids but not in cells transfected with all plasmids except pcDNA-PA (**Fig. 4B**). Similarly, a high luciferase activity was detected in the cells co-transfected with pHW2000-gLuc and pcDNA-PA, PB1, PB2, and NP plasmids, but nearly background levels were detected in cells transfected with pHW2000-Luc and pcDNA-PB1, PB2, and NP plasmids (**Fig. 4C**). Overall, there was a 10-fold increase in luciferase activity by the addition of the pcDNA-PA plasmid, suggesting a function of the RdRP activity from expression of PA, PB1, PB2, and NP.

**Fig. 4.**
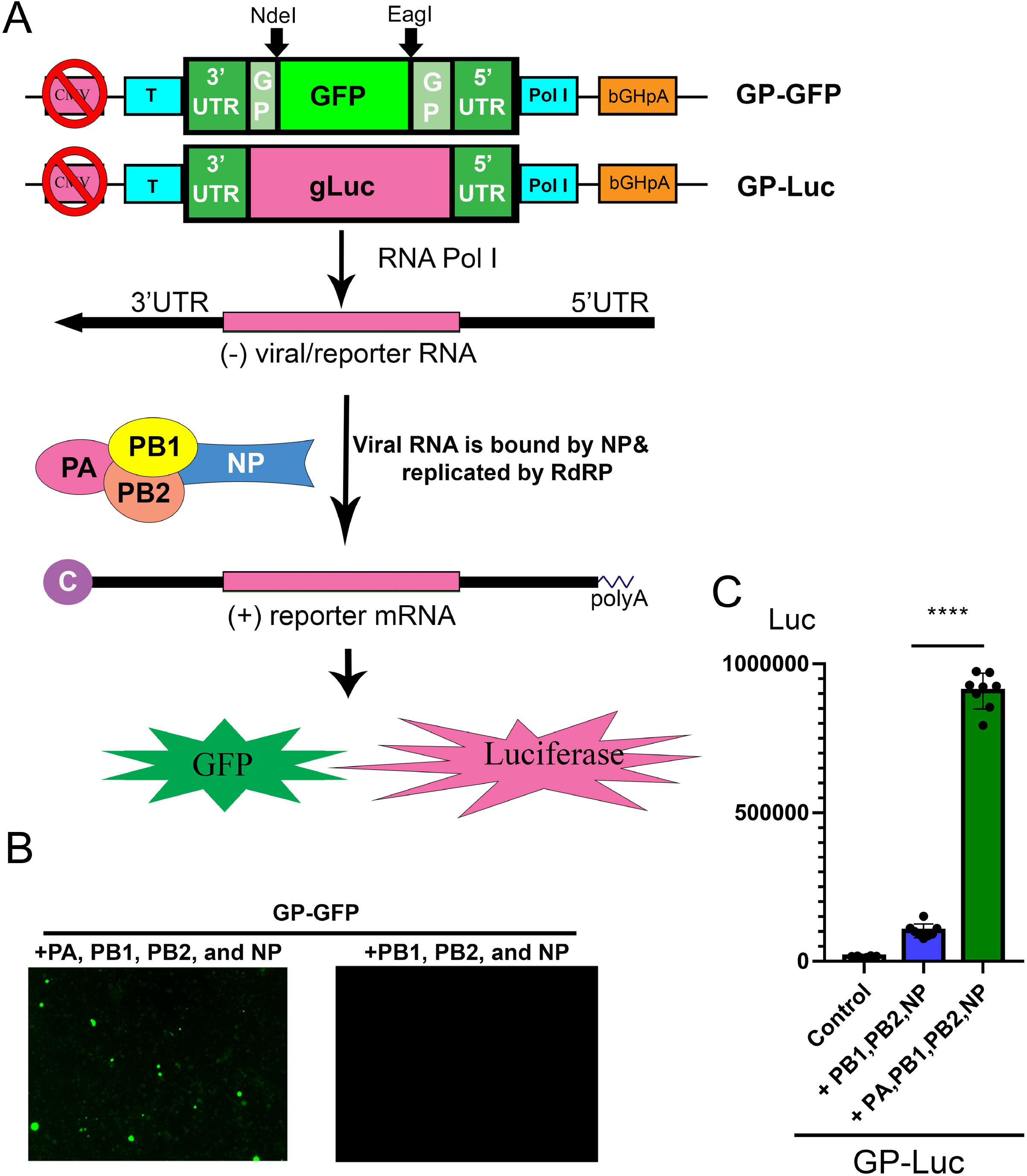
Establishment of a BRBV replicon report system. **(A) Diagram of the reporter system.** In a CMV pol II-deleted pHW-GP plasmid, a GFP open read frame (ORF) or a *Gaussia* luciferase (gLuc) ORF was inserted into the pol I expression cassette. In cells transfected with pcDNA-PA, PB1, PB2, and NP plasmids, pol l transcribes negative sense viral (reporter) RNA [(−)rRNA], which is used as a template for replication of (−)rRNA and production of reporter mRNA (GP-GFP or GP-gLuc) by the RdRP complex and NP protein as diagrammed. Then reporter mRNA is translated to express GFP or gLuc. Since the CMV pol II promoter was removed, the production of reporter mRNA is solely dependent on the replication of the reporter RNA (the activity of the RdRP complex). “C” at the 5’ end of the (+) reporter mRNA represents the 5’ m7G cap. **(B) Expression of PA, PB1, PB2, and NP supported GFP expression from the GP-GFP gene.** Co-transfection of the pHW-ΔCMV-GP-GFP with pcDNA-PA, PB1, PB2, and NP in HEK293T cells revealed green fluorescence under a UV microscope, but the co-transfection with only pcDNA-PB1, PB2, and NP did not. **(C) PA, PB1, PB2, and NP expression supported gLuc expression from GP-gLuc gene.** Co-transfection of the pHW-ΔCMV-GP-gLuc with pcDNA-PA, PB1, PB2, and NP in HEK293T cells showed a high luciferase activity (green bar), but not with only three pcDNA plasmids or no plasmids (control). All luciferase activities (Luc) were detected by the *Gaussia* Luciferase Glow Assay Kit (Pierce). Individual values are represented by black solid dots and the columns are the means with standard deviations from eight experiments performed in duplicate. ****, P < 0.0001 by one-way Student t-test.

### 3.4. Compound #9, a natural flavonoid myricetin, effectively, inhibits BRBV RdRP activity and BRBV replication in cells

We next used the BRBV replicon reporter system to test antiviral activity of three inhibitors of influenza virus RdRP, favipiravir, also known as T-705 (a pan RdRP inhibitor) (Furuta, et al., 2005; Fuchs, et al., 2019), pimodivir (VX-787), an inhibitor of PB2 cap-snatching activity (Byrn, et al., 2015; Furuta, et al., 2017), and baloxavir, a PA endonuclease inhibitor (Omoto, et al., 2018) (**Fig. 5A**). At 0.1, 1 and 10 μM, respectively, none inhibited luciferase (RdRP) activity by >50% (**Fig. 6A**). These drugs did not exhibit significant cytotoxicity at the tested concentrations in HEK293T cells (**Fig. 6C**).

**Fig. 5.**
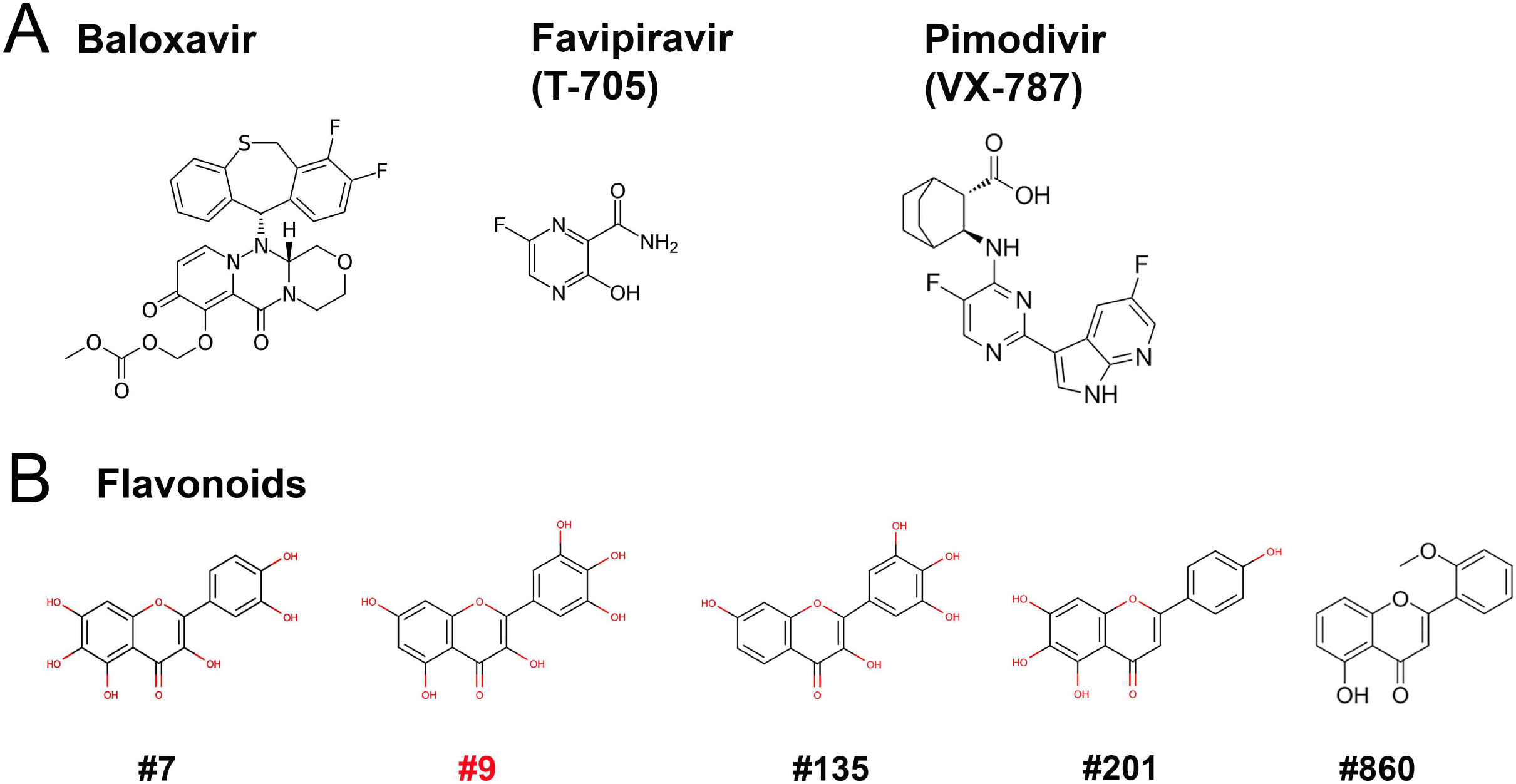
Compound structures. Structures of the three inhibitors of anti-RdRP of influenza (A) and 5 flavonoids (B) are shown.

**Fig. 6.**
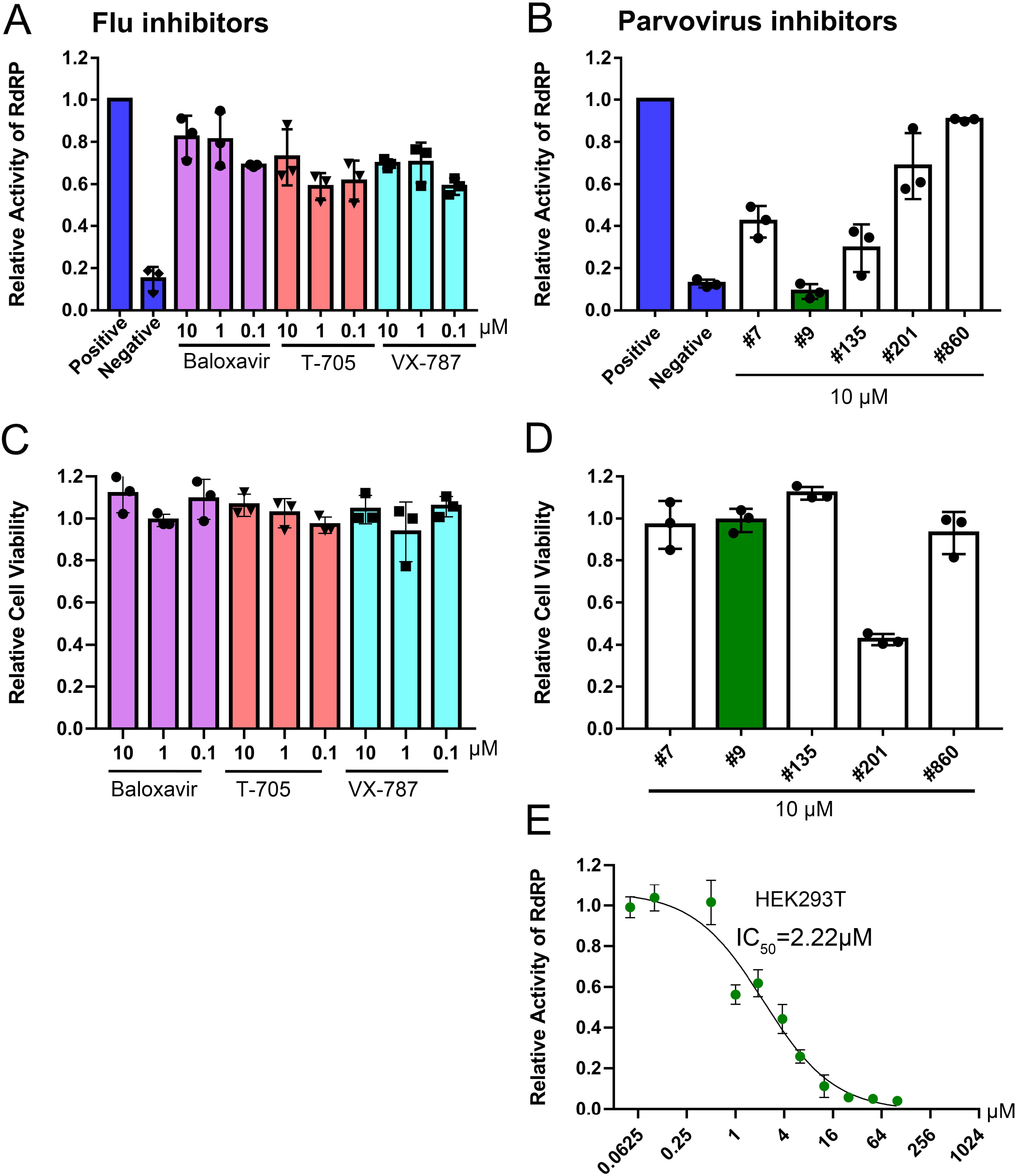
Screening of BRBV RdRP inhibitors using the BRBV replicon luciferase reporter. **(A&B) Inhibition of RdRP activity.** The inhibitory effect of different drugs, inhibitors targeting RdRP of influenza virus (Flu inhibitors) (A) and inhibitors of anti-parvovirus endonuclease compounds (Parvovirus inhibitors) (B), on BRBV RdRP activity was quantified using the GP-gLuc reporter assay at different concentrations as indicated. DMSO was used as a positive control of the reporter (Positive), and transfection with only pcDNA-PB1, PB2, and NP was used as a negative control of the reporter (Negative). At three days post transfection, the activity of the expressed *Gaussia* luciferase was quantified using a *Gaussia* Luciferase Glow Assay Kit (Pierce). Results from each compound were normalized to the vehicle control in each experiment and were expressed with the means and standard deviations obtained from three experiments performed in duplicate. **(C&D) Cytotoxicity assay**. Compounds were assayed for cytotoxicity in HEK293T cells at the concentrations indicated using a CytoTox-Glo™ Cytotoxicity Assay kit (Promega). The results are normalized to the mock-treated control (set as 1.0). Values are the means and standard deviations obtained from three experiments, each performed in duplicate. **(E) IC_50_ of compound #9 on RdRP activity.** Compound #9 was applied to HEK293T cells transfected with the gLuc reporter system of 5 plasmids at various concentrations as indicated. The polymerase activity was quantified by measuring the luciferase activity. Values are the means with standard deviations and are normalized to the mock group from three experiments performed in duplicate. The half maximal inhibitory concentration (IC_50_) was calculated with GraphPad Prism software.

Since influenza virus PA executes endonuclease activity on both single-stranded (ss)RNA and ssDNA (Dias, et al., 2009; Klumpp, et al., 2000; Doan, et al., 1999), we tested 4 flavonoid compounds (#7, #9, #135, and #201) (**Fig. 5B**) that inhibit nicking/endonuclease activity of the large nonstructural protein (NS1) of B19V on an ssDNA template (Xu, et al., 2018). Surprisingly, at 10 μM, 3 flavonoids (#7, #9, and #135) inhibited BRBV RdRP activity by >50%. In contrast, flavonoids #201 and #860 did not inhibit RdRP (**Fig. 6B**). Additional evidence of specific inhibition of the BRBV RdRP is provided by failure of the flavonoid #201 inhibited the endonuclease activity of B19V NS1 (Xu, et al., 2018). Although compound #201 was cytotoxic at 10 μM in HEK293T cells, compounds #7, #9 and #315 were not (**Fig. 6D**). The half maximal inhibitory concentration (IC_50_) of compound #9 in the reporter system was 2.22 μM (**Fig. 6E**). We next assayed the inhibitory effect of compound #9 on virus replication in both HEK293T and Vero cells. The half maximal effective concentration (EC_50_) of compound #9 against viral replication was 4.6 μM and 20.0 μM in HEK293T and Vero cells, respectively (**Fig. 6A&B**). Notably, compound #9 had negligible 50% cytotoxic concentrations (CC_50_) of 537 μM in HEK293T cells and >1 mM in Vero cells (**Fig. 7C&D**).

**Fig. 7.**
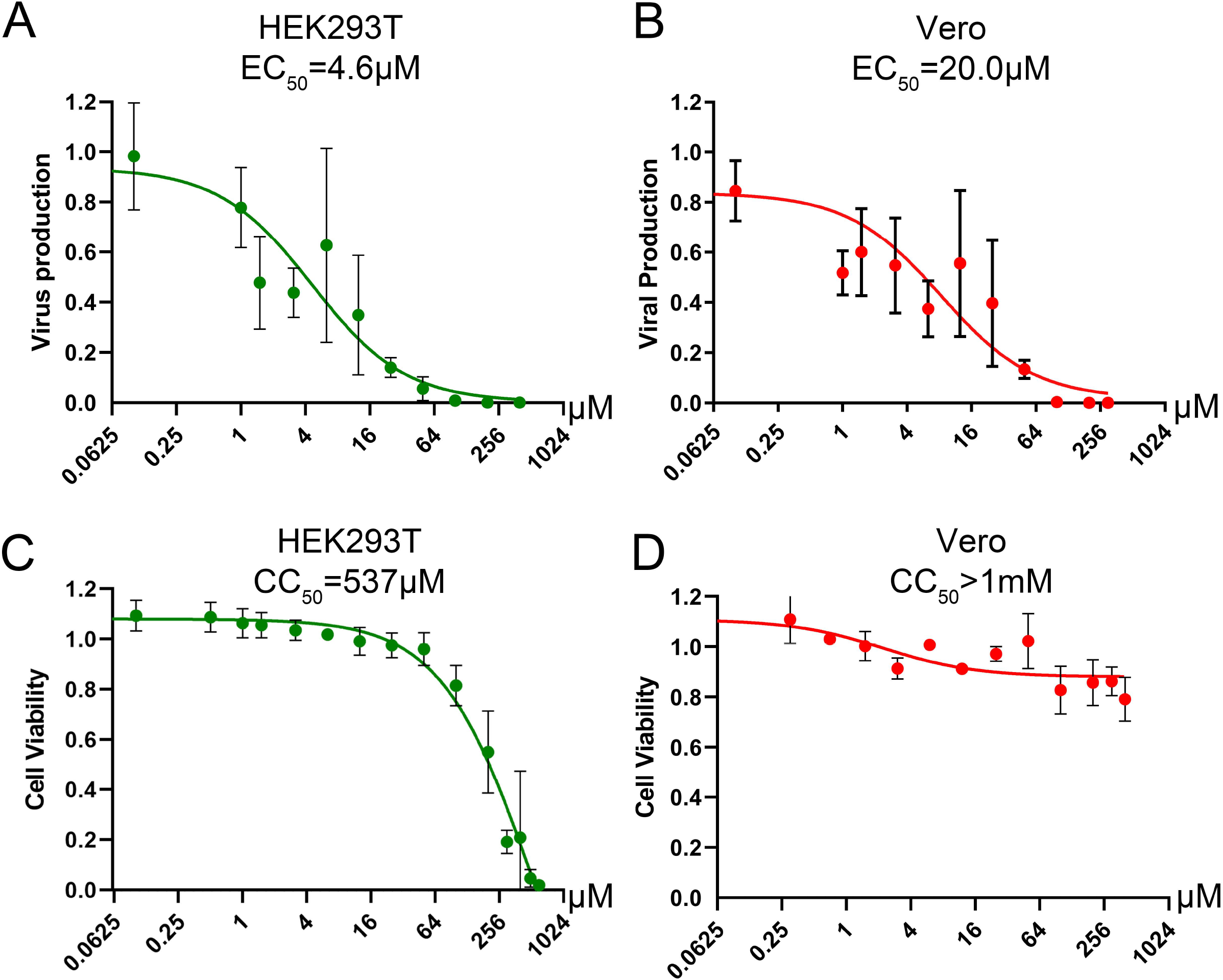
Determination of the CC_50_ and EC_50_ of compound #9 in cell cultures. **(A&B) EC_50_ of compound #9.** HEK293T cells (A) or Vero cells (B) were infected with BRBV at an MOI of 10, followed by application of compound #9 at various concentrations as indicated. At 3 days post treatment, RT-qPCR was performed to quantify the viral genome copy (vgc) numbers in the cell culture media. Values are the means with standard deviation and are normalized to a mock group, and were obtained from three experiments, each performed in duplicate. The EC_50_ was calculated using GraphPad Prism software. **(C&D) CC_50_ of compound #9**. A cytotoxicity assay was used to measure the viability of HEK293T cells (C) or Vero cells (D) affected by compound #9 at various concentrations as indicated. The percentage of viable cells were determined using a CytoTox-Glo cytotoxicity assay kit (Promega). The results are shown as relative values to the mock control cells. Values are the means with standard deviation obtained from three experiments performed in duplicate. The CC_50_ was calculated using GraphPad Prism software.

Collectively, our results demonstrated that a parvoviral endonuclease activity inhibitor, compound #9 (myricetin), inhibits BRBV replication in cells through inhibition of the RdRP activity with a therapeutic index (TI=CC_50_/EC_50_) of >50.

## 4. Discussion

In this study, we established a GFP/luciferase expression-based replicon reporter system for BRBV, and demonstrated it is suitable for screening of antivirals in a BSL2 setting. We identified myricetin (compound #9) as a potent anti-BRBV drug.

In contrast to influenza viruses, thogotoviruses are transmitted mainly through tick vectors and thus are also called “tick-borne viruses” (Haig, et al., 1965; Anderson & Casals, 1973). Phylogenetically, BRBV is most closely related to dhori virus and its subtype, batken virus, which have been known to occur in regions throughout Africa, Asia and Europe (Hubalek & Rudolf, 2012; Filipe, et al., 1985; Moore, et al., 1975; Lvov, et al., 1974). Dhori and batken viruses have been isolated from *Hyalomma* ticks, and antibodies against dhori virus have been detected in camels, goats, horses, cattle, and humans (Hubalek & Rudolf, 2012; Filipe, et al., 1985; Moore, et al., 1975; Lvov, et al., 1974). Interestingly, batken virus has also been isolated from several mosquito species (Lvov, et al., 1974). BRBV replicates at a high level in a variety of invertebrate and vertebrate cell lines, including HEK293T cells, and produces relatively high titers derived from all mammalian cells that are compatible with the known susceptibility of the human host to BRBV infection and diseases (Lambert, et al., 2015).

The cell cultured BRBV-KS has a series of variations in one α-helix of the NP protein, compared to the originally reported sequences (KP657749) by direct sequencing of the patient’s samples (**Fig. 3B**). We modeled the structure of BRBV NP protein for the amino acids’ electric charges and propensities to form an α-helix against the structure of influenza D virus (Donchet, et al., 2019) using SWISS-MODEL (Waterhouse, et al., 2018) (**Fig. 8**). The modeling indicates that the variations at aa 67-71 did not change the charges of any amino acids, had minimal effects on the polarity of the variant sequences, and the sequence was still able to form an α-helix. Interestingly, the BRBV-STL strain, which was isolated from a fatal BRBV case in St. Louis, Missouri, USA in 2017 (Bricker, et al., 2019), had the identical sequence at aa 67-71 in the NP protein (MK453525) as that of the cell cultured BRBV-KS.

**Fig. 8.**
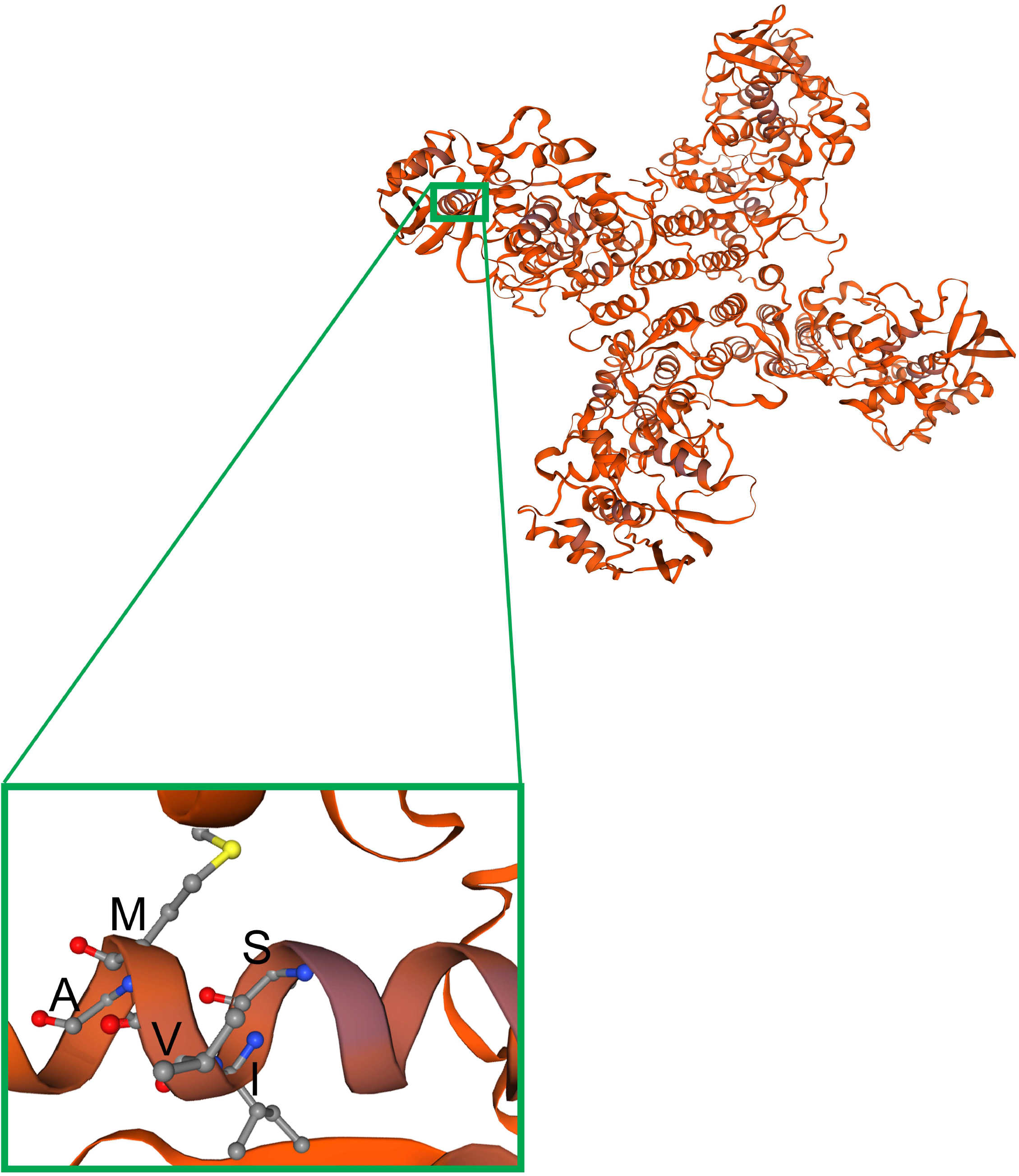
Adapted mutations in a α-helix structure of the NP protein. The BRBV NP structure was modeled on the template structure of the influenza D NP protein (Donchet et al., 2019) (SMTL ID: 5n2u.1) using SWISS-MODEL (Waterhouse, et al., 2018). NP forms a tetramer, as shown, which has four arms and a center to capture viral RNA and pass it to the RdRP. The region of NP carrying the α-helix that includes aa 67-71 (SIVMA) is enlarged from one of the monomers.

Currently, there is no treatment or vaccine for BRBV-infection caused diseases. Mice (CD-1) were susceptible to BRBV infection as evidenced by seroconversion, but the infection did not cause disease symptoms or death of the animals (Lambert, et al., 2015). Recently, mice lacking IFN-α/β receptor expression (IFNαAR^−/−^) have been used as an animal model to evaluate antivirals against BRBV (Bricker, et al., 2019; Fuchs, et al., 2019). Favipiravir, a pan inhibitor of RdRP (Furuta, et al., 2005), was shown to protect IFNαAR^−/−^ mice from lethal BRBV infection. The EC_50_ of favipiravir against BRBV infection of Vero was 310 μM (Bricker, et al., 2019). In animals, administration of 150 mg/kg of favipiravir twice daily at 3 days post-infection protected all animals from death of BRBV infection. However, it was speculated that favipiravir reached a concentration of 1.28 mM in serum, leading to the argument that it is not practical to use favipiravir to treat BRBV-infected patients. Notably myricetin, that was identified in our study, has a 15-fold lower EC_50_ (20 μM) compared with the 310 μM of favipiravir in Vero cells, suggesting that myricetin may be a more potent inhibitor of BRBV infection than favipiravir. Due to the lack of an animal BSL3 facility that can host BRBV-infected IFNαAR^−/−^ mice, we were not able to examine myricetin in animals.

Myricetin, a common plant-derived flavonoid, exhibits a wide range of activities, including antioxidant, anticancer, antidiabetic and anti-inflammatory activities, as well as antimicrobial activities (Semwal, et al., 2016). Myricetin was reported as a strong inhibitor of reverse transcriptase of retroviruses (Ono, et al., 1990). It has also been shown to be active in inhibition of nsp13, a helicase of the severe acute respiratory syndrome (SARS) coronavirus helicase that targets the ATPase activity in vitro (IC_50_=2.71 μM) (Yu, et al., 2012; Keum & Jeong, 2012). We found that myricetin does not exert obvious cytotoxicity in Vero and HEK239T cells, as well as in normal breast epithelial cells (Yu, et al., 2012; Keum & Jeong, 2012). We previously found that myricetin inhibited over 90% of the endonuclease activity of B19V NS1N (the endonuclease domain of NS1) in vitro at 10 μM (Xu, et al., 2018). We speculate that myricetin may inhibit the endonuclease activity of BRBV PA, which warrants further investigation.

## Acknowledgements

We are grateful to Dr. Zekun Wang and other members of the Qiu laboratory for technical support and valuable discussions. We thank Dr. Wenjun Ma at Kansas State University for providing the pHW2000 plasmid. The following reagent was obtained through BEI Resources, NIAID, NIH: Bourbon Virus, Original, NR-50132.

This study was supported by PHS grant AI144564 from the National Institute of Allergy and Infectious Diseases. The funders had no role in study design, data collection and interpretation, or the decision to submit the work for publication.

